# Ketone ester-enriched diet ameliorates motor and dopamine release deficits in MitoPark mice

**DOI:** 10.1101/2022.11.14.516368

**Authors:** Vikrant R. Mahajan, Jacob A. Nadel, M. Todd King, Robert J. Pawlosky, Margaret I. Davis, Richard L. Veech, David M. Lovinger, Armando G. Salinas

## Abstract

**Background:** Parkinson’s disease is a progressive, neurodegenerative disease characterized by motor dysfunction and dopamine deficits. The MitoPark mouse recapitulates several facets of Parkinson’s disease, including gradual development of motor deficits, which enables the study of potential therapeutic interventions. One therapeutic strategy involves decreasing the mitochondrial metabolic load by inducing ketosis and providing an alternative energy source for neurons, leading to decreased neuronal oxidative stress.

**Objective:** We assessed whether administration of a ketone ester-enriched diet would improve motor and dopamine release deficits in MitoPark mice.

**Methods:** Motor function (rotarod and open field tests), dopamine release (fast-scan cyclic voltammetry), tissue dopamine levels (GC-MS), and dopamine neurons and axons (immunofluorescence) were assessed in MitoPark and control mice fed either the standard or ketone ester-enriched diets.

**Results:** When started on the ketone diet before motor dysfunction onset, MitoPark mice had preserved motor function relative to standard diet MitoPark mice. While the ketone ester enriched diet did not preserve dopamine neurons or striatal dopamine axons, dopamine release in ketone diet MitoPark mice was greater than standard diet MitoPark mice but less than control mice. In a follow up experiment, we began the ketone diet after motor dysfunction onset and observed a modest preservation of motor function in ketone diet MitoPark mice relative to standard diet MitoPark mice.

**Conclusion:** The improvement in motor dysfunction indicates that a ketone ester enriched diet or ketone supplement may represent a promising adjunct treatment for Parkinson’s disease.

## Introduction

Parkinson’s disease (PD) is a progressive, neurodegenerative disease characterized by hypokinesia, resting tremor, and dystonia.^1^ It is the second most common neurodegenerative disease with a 1% prevalence in the population over 60 years old.^2^ The underlying pathology of PD involves a progressive loss of dopaminergic neurons in the substantia nigra pars compacta (SNc),^1^ however, the exact mechanisms underlying this loss remain elusive. Converging lines of evidence suggest a central role for mitochondrial dysfunction in PD etiology.^3^ Indeed, toxins targeting mitochondrial respiration produce selective vulnerability in SNc dopamine neurons,^4, 5^ and PD-like behavioral disturbances. Additionally, dopamine neurons from PD patients were found to have more mitochondrial DNA deletions than age-matched controls.^6^ Specifically, mitochondrial dysfunction is hypothesized to increase oxidative stress.^7^ The MitoPark (MP) model is a mouse model of PD in which the mitochondrial transcription factor TFAM A is selectively knocked out of dopamine neurons, resulting in a progressive dopaminergic neuron degeneration and a gradual onset of PD-like motor deficits.^8^ This model also recapitulates several important aspects of PD pathophysiology, such as the gradual development of motor deficits in adulthood, responsiveness to L-DOPA, and protein inclusions.^9, 10^ Recent work has attributed degeneration in MitoPark mice to impaired antioxidant defense, which is itself due to mitochondrial dysfunction and calcium dishomeostasis.^11^ Thus, the MP model is suitable not only to capture the hallmark progressive pathophysiology of PD but also to study the effects of potential therapeutic interventions.

In PD, L-DOPA provides effective temporary motor symptom relief but does not address the underlying pathology of the disorder. Furthermore, neither L-DOPA nor other dopamine replacement therapies help to preserve dopamine neuron function. The high-fat/low-carbohydrate (ketogenic) diet has been examined recently for its potential application in ameliorating neurological disease pathologies by inducing ketosis.^12-14^ Exogenous oral administration of D-β-Hydroxybutyrate (D-BHB), in the form of a ketone ester enriched diet (KEED), has been shown to ameliorate symptoms in rodent models of Alzheimer’s,^15^ metastatic cancer,^16^ diabetes,^17^ and epilepsy.^18, 19^ Furthermore, the bioenergetics of D-BHB gives a mechanism of action for sequestering and quenching of reactive oxygen species.^20^ Given the aforementioned roles of mitochondrial dysfunction and oxidative stress in PD, we hypothesized that the KEED would alleviate PD symptoms in MitoPark mice, potentially by reducing dopamine neuron metabolic load and/or increasing antioxidant availability.

Thus, we utilized the MitoPark mouse model to assess the therapeutic potential of a KEED to curtail PD-like symptoms. Specifically, we assessed the progressive loss of motor function under normal (open field) and challenging (rotarod) conditions in MitoPark and control mice fed a standard diet (SD) or a KEED. In our first experiment, we began mice on the KEED before symptom onset and observed a significant preservation of motor function in MitoPark mice fed the KEED. This was paralleled by a modest preservation in dorsal striatal dopamine release measured with fast-scan cyclic voltammetry (FSCV). Interestingly, this preservation of striatal dopamine release was not correlated with preserved dopamine neurons or dorsal striatal dopamine axon density in MitoPark mice fed the KEED. Similarly, the KEED did not increase or preserve tonic dopamine (or its primary metabolite DOPAC) levels in dorsal striatum of MitoPark mice. In a follow up experiment, we began feeding mice with the KEED after the onset of motor dysfunction and observed a more modest rescue of locomotor activity in the rotarod, but not open field, assay. Altogether, our results suggest that a KEED may be a promising adjunct therapeutic treatment for PD.

## Materials and Methods

For full details see Supplemental Materials.

### Animals

All procedures were performed in compliance with the National Institutes of Health Care and Use of Animals guidelines and approved by the Institutional Animal Care and Use Committee of the National Institute on Alcohol Abuse and Alcoholism. MitoPark and control mice were generated as previously described.^8, 9, 21^ Mice were housed up to four per cage on a 12 h light/dark cycle in a temperature- and humidity-controlled room with ad libitum access to food and water.

### Ketone ester enriched diet

The KEED has been described previously.^22^ Briefly, standard rodent diet was modified to include a ketone ester (D-β-hydroxybutyrate and R-1,3-butanediol) at the caloric expense of carbohydrates (Figure 1a). We tested diets with 16%, 33%, and 50% of calories from ketone esters and chose the 16% KEED for use in subsequent experiments.

**Figure 1.**
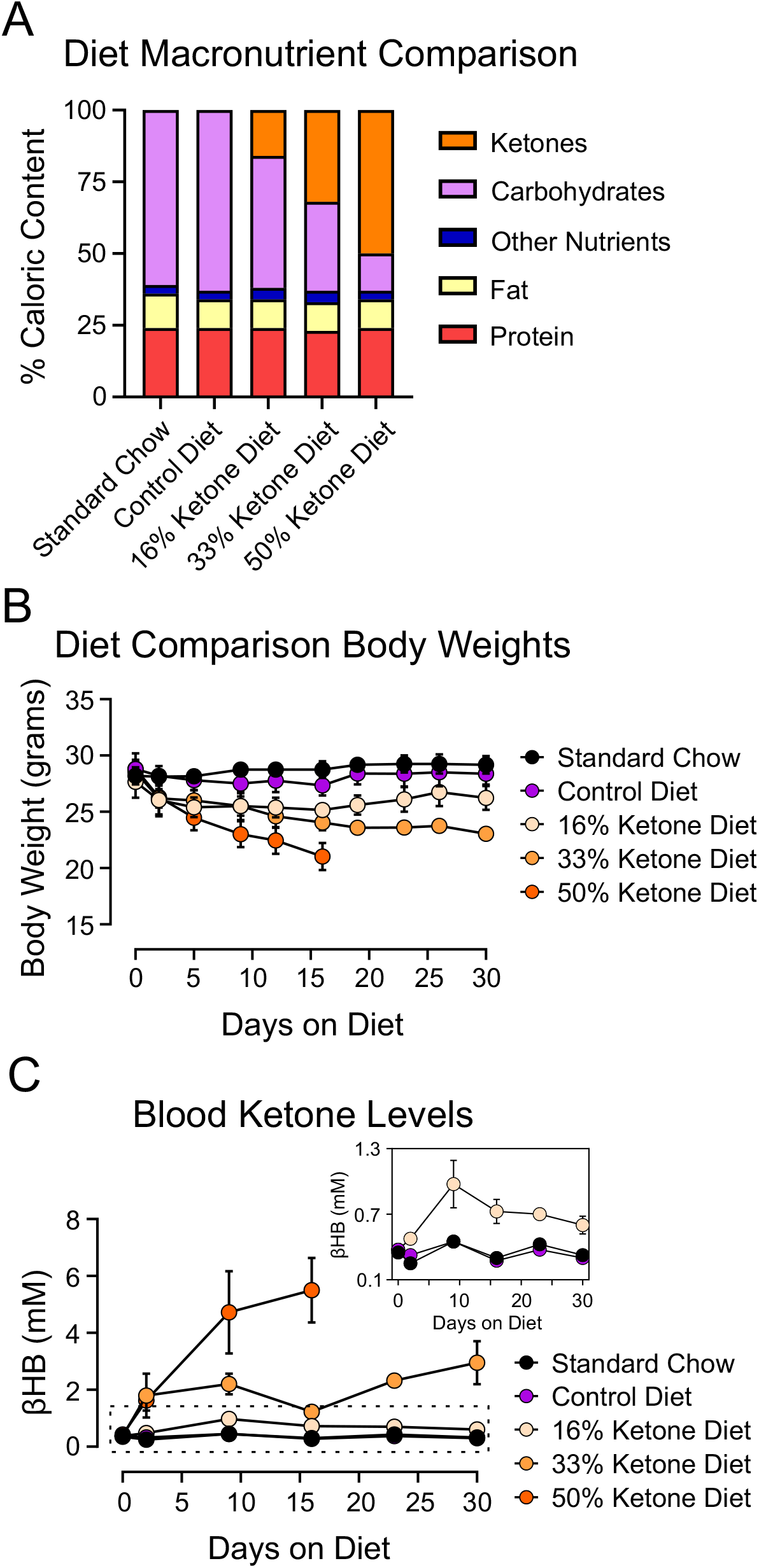
Comparison of ketone ester enriched diets (KEED). (A) Macronutrient composition of tested KEEDs. Mice were fed each KEED for 30 days with body weights (B) and blood ketone levels (C) assessed periodically. n=4 mice per diet group.

### Blood ketone levels and diet testing

Male C57BL/6J mice (Jackson Laboratory, strain 000664) were used to determine achievable blood ketone levels for each diet: standard diet, control diet, 16% KEED, 33% KEED, and 50% KEED. Tail vein blood samples were collected at 0, 2, 9, 16, 23, and 30 days after starting the KEED. β-hydroxybutyrate levels were measured using Precision-Xtra blood ketone test strips and a Precision-Xtra meter (Abbott Labs, Abbott Park, IL, USA). Due to the circadian fluctuation of ketone levels,^23, 24^ blood samples were obtained at the same point in the light cycle for each experiment.

### Accelerating rotarod and open field

At four weeks of age, subjects were assigned to the standard diet (SD) or 16% KEED groups. Beginning at six weeks of age, MitoPark and control mice fed either the SD or KEED were run in an open field task to assess spontaneous movement. Each week, mice were placed in large cages (42cm x 22.5cm x 25cm) and video recorded for one hour. The videos were processed in EthoVision (Noldus, Leesburg, VA) and average velocity for each mouse/session was calculated. On a different day in the same week, mice were run on an accelerating rotarod (EZRod, Omnitech Electronics or MedAssociates) to assess motor coordination and challenged movement. Mice were run on the same lane and rotarod for the duration of testing. Mice were tested for 5 trials/day with the rotarod speed increasing from 4 to 40 rpm over 300s. Trials were conducted every 10 min. The latency to fall from the rotarod was recorded for each trial. Individual trials were stopped and latency was recorded if mice held onto the rod for two consecutive rotations or reached 40 rpm. Beginning at eight weeks of age, a second group of mice was tested every other week on the rotarod and in the open field. These mice began the KEED at 14 weeks, after the onset of significant motor deficits.

### Fast-scan cyclic voltammetry

At 5, 10, or 20 weeks of age, mice were used for FSCV experiments to assess changes in dorsal striatal dopamine release. Briefly, 300μm-thick brain slices containing the dorsal striatum were prepared as previously described.^25^ Dopamine release was electrically evoked using a range of stimulation intensities with a DS3 constant current stimulus generator (Digitimer NA, Ft. Lauderdale, FL) and a twisted, stainless steel stimulating electrode (P1 Technologies, Roanoke, VA). Carbon fiber electrodes were made as previously described^26, 27^ and calibrated post hoc against a 1μM dopamine solution standard. All KEED mice used for these experiments began the diet at 4 weeks.

### Immunofluorescence

Mice were anesthetized and transcardially perfused with PBS followed by a 4% paraformaldehyde/PBS solution. Brains were post-fixed overnight before sectioning into 50μm thick sections. The sections were washed 4×10 minutes in PBS-T (0.2% Triton X-100), then 2×10 minutes in 0.5% NaBH4 in PBS, then 4×10 minutes in PBS-T before blocking for two hours in 5% BSA in PBS-T. Sections were then incubated overnight at 4°C in a primary antibody/0.5% BSA/PBS-T solution of rabbit anti-TH (1:1000, Invitrogen, Carlsbad, CA, 701949). Sections were then washed 6×10 minutes in PBS-T before incubating in secondary antibody/0.5% BSA/PBS-T solution of Alexa568 donkey anti-rabbit (1:2000) for two hours. Sections were washed 6×10 minutes with PBS, mounted onto subbed slides, cover-slipped with DAPI Fluoromount-G (Electron Microscopy Sciences, Hatfield, PA), and imaged with a Zeiss AxioZoom microscope with ZEN software. Images were quantified using Fiji/Image J by a researcher blinded to the treatment and genotype of the mice. All KEED mice used for these experiments began the diet at 4 weeks.

### Biochemistry

Dorsal striatal tissues from 20-week-old MitoPark and control mice fed the SD or KEED were obtained and processed for dopamine and DOPAC levels with gas chromatography-mass spectrometry (GC-MS). Analytes were analyzed as its tertiary butyl dimethylsilyl ether-ester derivatives using GC-MS in the electron impact mode and quantified using the ^2^H_3_-acetate. Briefly, samples were homogenized, centrifuged, and the sample supernatant was collected and dried under nitrogen along with labeled internal standards before sylilation and processing on an Agilent 5973 quadrupole GC-MS (Agilent, Wilmington, DE). The ratios of the ions m/z 354/358 (dopamine retention time (RT), 13.1 min) and 384/389 (DOPAC, RT 12.85 min) were used to determine the quantity of each analyte.

### Reagents

Unless otherwise indicated, reagents were obtained from Sigma-Aldrich (St. Louis, MO). AlexaFluor-conjugated secondary antibodies were obtained from Invitrogen (Carlsbad, CA).

### Statistics

GraphPad Prism 8 was used for graphing and statistics. Where appropriate, we utilized one-, two-, and three-way ANOVAs (with repeated measures and Geisser-Greenhouse corrections where necessary), as well as mixed-effects analysis for data sets with missing values. When three-way ANOVAs or mixed effects analyses had significant interactions, we performed two-way analyses on the appropriate subgroups to better understand the differences between groups. Multiple comparisons were performed using Tukey’s or Sidak’s multiple comparisons tests.

## Results

For detailed results, see supplemental materials.

### Comparison of ketone ester enriched diets

We compared KEEDs with 16%, 33%, and 50% of carbohydrate calories replaced by D-BHB, alongside standard rodent chow and a control diet (Fig 1A) to maximize blood ketone levels with KEED palatability. We found that all KEEDs elevated blood ketone levels and that mice on each KEED initially lost body weight when switched to their respective diet (Fig 1B). Mice on the 16% KEED regained body weight by the third week on the diet, whereas mice on the 33% KEED did not recover their initial body weights by 30 days. Mice fed the 50% KEED were removed from the study at 16 days on the KEED due to severe weight loss. Given the weight loss in the 33% and 50% KEED groups, we chose the 16% KEED for subsequent experiments.

### KEED ameliorates spontaneous locomotor deficits

From 6-20 weeks, mice were tested weekly in the open field (spontaneous locomotion) and on an accelerating rotarod (forced locomotion). To decrease individual week-to-week performance variability, behavioral data was averaged into epochs: Early (weeks 6-10), Mid (11-15), and Late (16-20). In the open field, mixed-effects three-factor mixed modeling analyses indicated significant effects of week (F_(2,123)_=8.097, p=0.0005) and genotype (F_(1,64)_=15.92, p=0.0002), as well as significant week x genotype (F_(2,123)_=20.76, p<0.0001) and genotype x diet (F_(1,64)_=10.61 p=0.0006) interactions, and a significant week x genotype x diet interaction (F_(2,123)_=4.291, p=0.0158). To investigate simple main effects, we performed two-way mixed-effects modeling across genotype and diet factors.

#### Diet

In control mice, there were significant effects of week (F_(1.457,48.82)_=4.591, p=0.0242), and diet (F_(1,35)_=5.081, p=0.0306). The significant effect of week was driven by an increased average velocity between Early and Mid epochs (p=0.0162). The significant effect of diet was driven by the C-k mice having a significantly higher average velocity than C+k mice in the Late epoch (p=0.0370). In MitoPark mice, there were significant effects of week (F_(1.762,49.33)_=31.78, p<0.0001) and diet (F_(1,29)_=5.872, p=0.0219), and a trend towards a significant interaction (F_(2,56)_=3.134, p=0.0513). The effect of week was driven by a decrease in average velocity across epochs (p<0.002 for all). The effect of diet was driven by the MP+k mice moving significantly more than the MP-k mice in Late weeks (p=0.0012). Furthermore, although MP-k mice significantly decreased average velocity across epochs (Early vs. Mid, p=0.0129; Mid vs. Late, p=0.0002; Early vs. Late, p<0.0001), MP+k mice only showed significant deficits when comparing Early and Late epochs (Early vs. Mid, p=0.1322, Mid vs. Late, p=0.4888, Early vs. Late, p=0.0004). Combined, these data suggest that MitoPark mice fed a ketone diet develop delayed deficits in spontaneous locomotion relative to MitoPark mice fed SD.

#### Genotype

In mice fed SD, there was a significant effect of week (F_(1.435,48.80)_=4.074, p<0.0351) and genotype (F_(1,36)_=22.62, p<0.0001), and a significant interaction (F_(2,68)_=16.70, p<0.0001). C-k mice displayed slower average velocity only on Early vs. Mid (p=0.0475), but not Early vs. Late or Mid vs. Late epochs. In contrast, MP-k mice progressively declined across time points (Early vs. Mid, p=0.0129; Mid vs. Late, p=0.0002, Early vs. Late, p<0.0001). Furthermore, MP-k mice had significant spontaneous movement deficits at both Mid (p<0.0001) and Late (p<0.0001) time points, compared to C-k. In KEED fed mice, there was a significant effect of week (F_(1.605,44.14)_=7.074, p=0.0040) and a week x genotype interaction (F_(2,55)_=7.045, p=0.0019). The effect of week was driven by the MP+k mice, whose velocity declined between Early and Late epochs (p=0.0004). However, C+k and MP+k mice did not differ at any timepoint (p<0.05). Overall, the presence of distinct effects across genotypes in SD but not KEED mice supports a beneficial effect of the KEED.

### KEED ameliorates forced locomotion deficits on rotarod

To assess forced locomotion, motor coordination, and balance, mice were run weekly on an accelerating rotarod. Mixed-effects three-factor mixed modeling analyses revealed significant effects of week (F_(2,113)_=59.53, p<0.0001) and genotype (F_(1,59)_=12.35, p=0.0009). Furthermore, there were significant week x genotype (F_(2,113)_=18.64, p<0.0001) and week x diet (F_(2,113)_=0.5810, p=0.0040) interactions, and a significant week x genotype x diet interaction (F_(2,113)_=4.956, p=0.0086). To investigate simple main effects, we performed two-way mixed-effects modeling, with mice split by genotype or diet factors.

#### Diet

In control mice, there was a significant effect of week (F_(1.863,54.96)_=5.745, p=0.0064). Tukey’s multiple comparisons tests indicated the significant effect of week was driven by a decreased latency to fall at the later weeks relative to early weeks (p=0.0098) across diets. In MitoPark mice, there was a significant effect of week (F_(1.895,51.16)_=75.88, p<0.0001) and a significant week x diet interaction (F_(2,54)_=11.22, p<0.0001), but no significant effect of diet alone (F_(1,29)_=3.3702, p=0.0642). MitoPark mice showed progressive decline across time (MP-k: all comparisons p<0.0001; MP+k: Early vs. Mid, p=0.0264, Mid vs. Late, p=0.0166, Early vs. Late, p=0.0007). However, MP+k mice had significantly longer latency to fall in Late weeks compared to the MP-k mice (p=0.0014), indicating the KEED ameliorated forced locomotor deficits. Overall, these data indicate the KEED may improve forced locomotion in MitoPark mice.

#### Genotype

In SD mice, there was a significant effect of week (F_(1.886, 55.65)_=50.10, p<0.0001) and genotype (F_(1,31)_=12.18, p=0.0015), and a significant interaction (F_(2,69)_=21.05, p<0.0001). Multiple comparisons tests indicated that C-k mice did not change performance across weeks, but in MP-k mice, latency to fall decreased across epochs (all p<0.0001). Although latency to fall in MP-k mice declined across epochs, these mice only differed from the C-k mice at Late weeks (p<0.0001). In KEED mice, there was a significant effect of week (F_(1.889,51.01)_=14.79, p<0.0001) but no genotype or interaction effects. Tukey’s multiple comparisons tests indicate that the effect of week is driven by significant, or near-significant, differences between all time points (Early vs. Mid, p=0.0081; Mid vs. Late, p=0.0580; Early vs. Late, p=0.0003) regardless of diet. These analyses also support the ameliorative effect of the KEED on forced movement and motor coordination.

### KEED ameliorates dopamine release deficits

We assessed evoked dopamine release using FSCV in striatal slices from control and MitoPark mice fed control diet at 5, 10, and 20 weeks of age. We also assessed dopamine release in 20-week-old mice fed the KEED. At 5 weeks, control and MitoPark mice did not differ in evoked dopamine release. At 10 weeks, MitoPark mice had lower dopamine release overall (significant main effect of genotype, F_(1,18)_=8.902, p=0.0080; significant main effect of intensity, F_(1.185,21.33)_=22.44, p<0.0001; significant interaction, F_(5,90)_=3.032, p=0.0142). At 20 weeks, three-way ANOVA revealed significant effects of stimulation intensity (F_(1.524,47.23)_=55.02, p<0.0001), genotype (F_(1,31)_=57.27, p<0.0001), and a significant genotype x intensity interaction (F_(5,155)_=37.97, p<0.0001). Simple main effects analysis indicated that dopamine release in C-k and C+k mice did not differ. However, dopamine release significantly differed between MP-k and MP+k groups (significant effect of diet (F_(1,15)_=18.74, p=0.0006) and intensity x diet interaction (F_(5,75)_=19.43, p<0.0001)). Post-hoc tests indicated these effects were driven by significant or near-significant increases in dopamine release in the MP+k mice at 600 (p=0.0483) and 800 (p=0.0579) μA.

### KEED does not increase cell survival, striatal TH+ axons, or tissue dopamine content

To determine if the KEED rescue of dopamine release was due to preservation of dopamine neurons or axons, we performed immunohistochemistry for tyrosine hydroxylase (TH) (Fig 4A). Across the anterior-posterior gradient of the dorsal striatum, MitoPark mice had significantly decreased TH+ immunoreactivity compared to controls (three-way repeated measures mixed-effects analysis, significant main effect of genotype, F_(1,12)_=435.0, p<0.0001), indicating dopamine axons degenerated regardless of diet (Fig 4B). Similarly, TH+ cell counts in the SNc and VTA indicated a genotype-but not diet-dependent decrease in cell count (three-way repeated measures ANOVA, significant main effect of genotype, F_(1,12)_=19.60, p=0.0008) (Fig 4C). GC-MS analyses of flash-frozen striatal tissue indicated that, although dopamine and DOPAC content was significantly diminished in MitoPark mice (two-way ANOVAs, dopamine: significant main effect of genotype, F_(1,31)_=245.1, p<0.0001; DOPAC: significant main effect of genotype, F_(1,31)_=81.38, p<0.0001), the KEED did not increase content of either compound (Fig 4D-E). Sidak’s multiple comparisons tests confirmed that MitoPark mice, regardless of diet, had similar dopamine and DOPAC levels. Groups did not differ in DOPAC/DA ratio, which is thought to be a correlate of MAO activity and levels of oxidative stress.^28^ Thus, despite the beneficial KEED effects on motor deficits and dopamine release, the diet does not seem to slow neurodegeneration or increase striatal dopamine levels.

### Late KEED intervention delays but does not rescue motor deficits

Our initial experiment prevented the development of motor deficits in MitoPark mice up to 20 weeks. However, we began the KEED at 4 weeks, before any motor deficits emerged. This schedule is not comparable to treatment of PD in humans, as treatments begin after symptom onset. Thus, we tested whether the KEED would benefit subjects after the onset of motor deficits. Mice began rotarod and open field testing at 8 weeks and KEED began at 14 weeks (after onset of open field and rotarod deficits in MP-k mice, Fig 2). The endpoint was extended to 22 weeks to better capture the potential effects of KEED intervention (Fig 5A). Behavioral data were binned into epochs as before (8-10, 12-14, 16-18, and 20-22 weeks).

**Figure 2.**
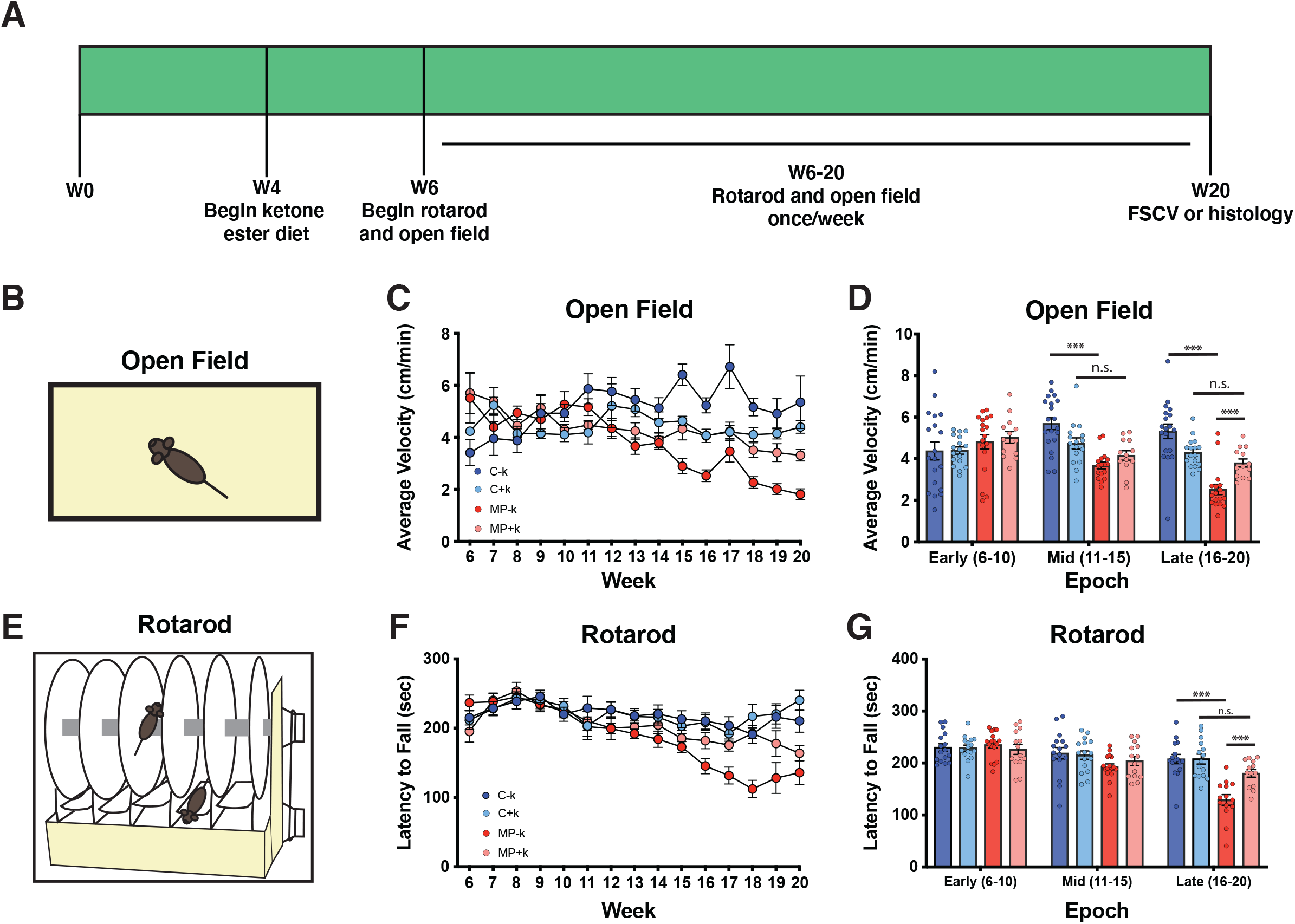
Presymptomatic KEED treatment ameliorated motor deficits in MitoPark mice. (A) Experimental timeline illustrating behavioral testing and diet onset. (B-D) Mice were tested in an open field to assess locomotion. MitoPark mice fed the SD exhibited decreased locomotion compared to MitoPark mice fed the 16% KEED (D). Mice were further tested in the rotarod apparatus (E). MitoPark mice fed the SD performed progressively worse (F) than KEED MitoPark and control mice (G). n=10-14 mice per group. *** p<0.001; n.s. p>0.1

**Figure 3.**
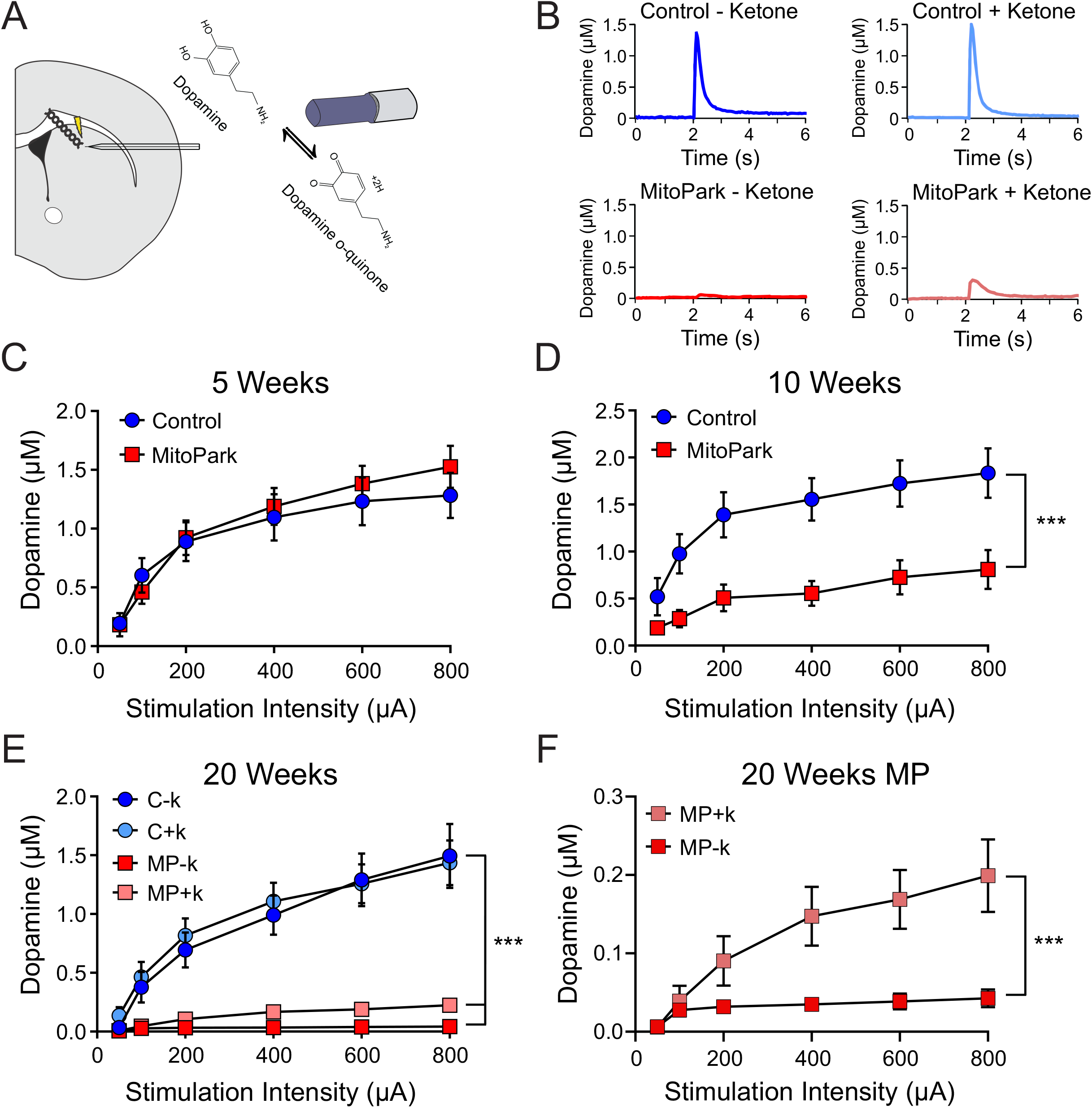
Progressive decrease in dopamine release in MitoPark mice is ameliorated by KEED. (A) Schematic for fast-scan cyclic voltammetry experiments. (B) Representative dopamine release data for each group at 20 weeks. At 5 weeks (C), dopamine release does not differ between genotypes. (D) At 10 weeks, dopamine release is decreased in MitoPark mice. (E) At 20 weeks, dopamine release was decreased in MitoPark relative to control mice in SD groups. (F) MitoPark mice fed the KEED had greater dopamine release than MitoPark mice fed the SD. n=7-12 slices per group from >5 mice per group. *** p<0.001

**Figure 4.**
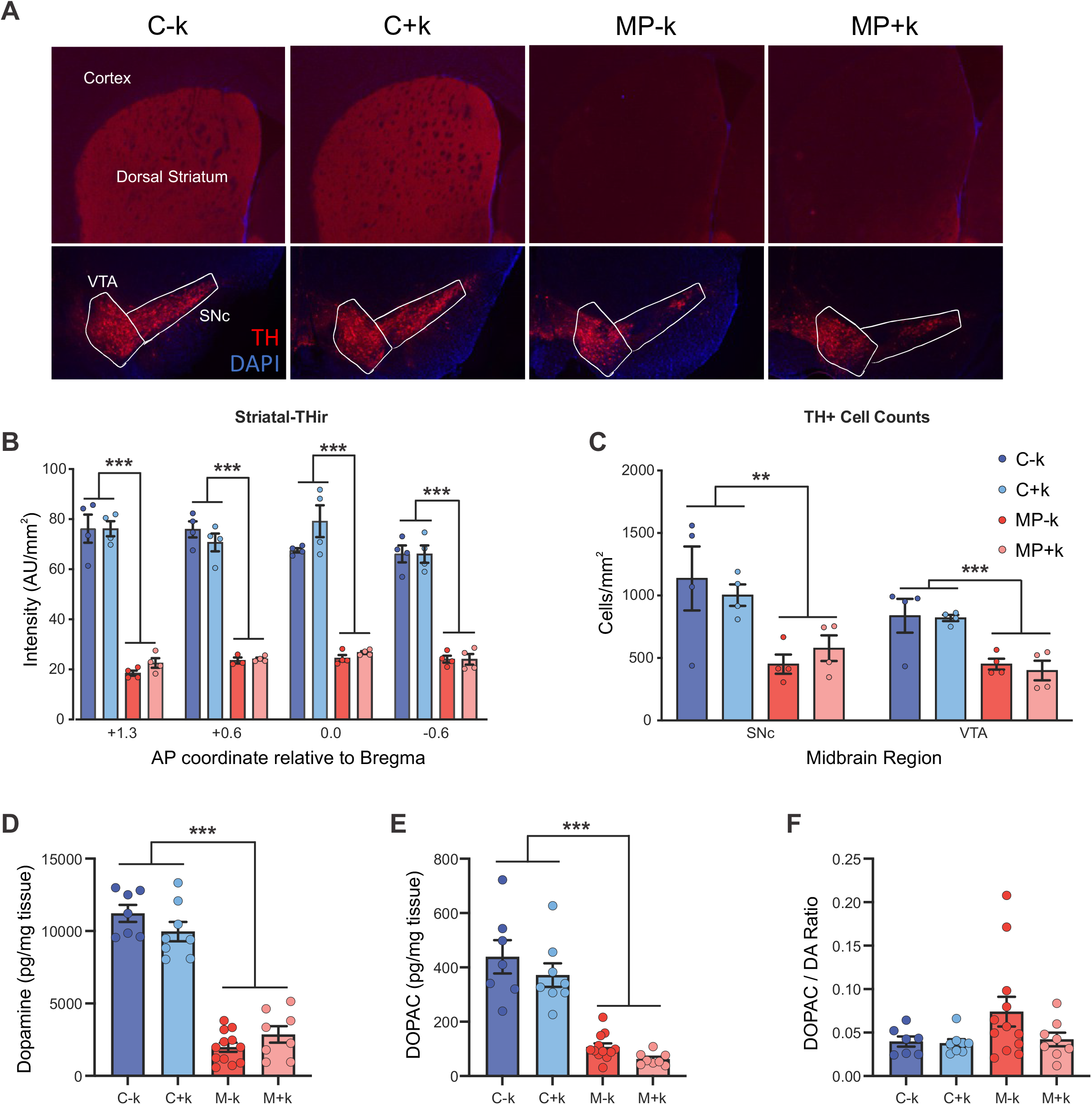
KEED does not preserve dopamine neurons or striatal axon loss in 20-week-old MitoPark mice. (A) Representative immunohistochemistry images of dorsal striatum (upper row) and midbrain dopamine neurons (lower row). (B) Striatal tyrosine hydroxylase-immunoreactivity and midbrain dopamine neurons (C) were decreased in both MitoPark groups. Dopamine (D) and DOPAC (E) content were decreased in dorsal striatal tissue from 20-week-old MitoPark mice relative to controls. DOPAC/Dopamine ratios (F) did not differ significantly across groups. n=4 mice per group for immunohistochemistry experiments and n=7-12 mice per group for biochemistry experiments. ** p<0.01; *** p<0.001

**Figure 5.**
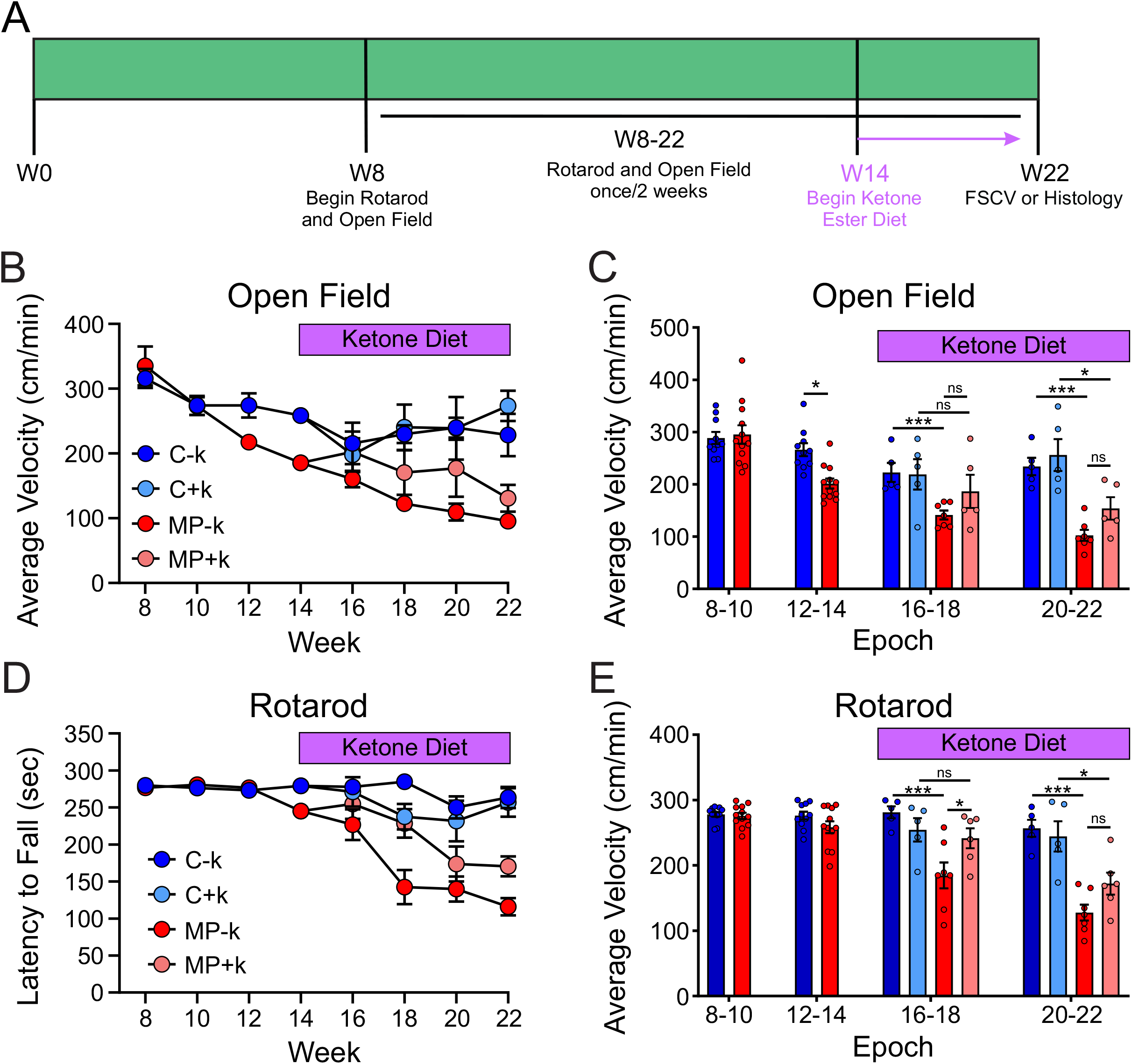
Beginning ketone ester enriched diet after motor deficit onset may slow progressive motor dysfunction in MitoPark mice. (A) Experimental timeline illustrating behavioral testing and KEED onset. (B&C) MitoPark mice fed either diet exhibited decreased locomotion in the open field relative to control groups. (D&E) MitoPark mice fed the control diet performed progressively worse on the rotarod across the study (D). MitoPark mice fed the KEED had improved motor function initially but failed to maintain this improvement. n=5-7 mice per group. * p<0.05; *** p<0.001

Before KEED intervention (first two epochs) in the open field, we found a significant effect of time (two-way RM ANOVA, F_(1,20)_=47.95, p<0.0001) and a significant time x genotype interaction (F_(1,20)_=18.12, p=0.0004), but no effect of genotype alone. Sidak’s multiple comparisons tests indicated that MitoPark, but not control, mice decreased movement from early to mid-time points (p<0.0001), and that at the mid time point, MitoPark mice moved significantly less than controls (p=0.0031). After dietary intervention (final two epochs), a significant effect of genotype (three-way RM ANOVA, F_(1,18)_=20.24, p=0.0003), and time x genotype interaction (F_(1,18)_=14.03, p=0.0015) were observed. All other effects and interactions did not reach statistical significance (p>0.05). To further investigate the effects that reached significance, we analyzed the data across genotype and diet factors with two-way RM ANOVAs.

#### Genotype

For SD mice (C-k and MP-k), there were genotype (F_(1,10)_=38.68, p<0.0001) and time x genotype interaction (F_(1,10)_=12.94, p=0.0049) effects. Sidak’s multiple comparisons tests revealed that this effect was due to MP-k mice moving less over time (p=0.0031), and MitoPark mice moving less at both post-diet time points (16-18: p=0.0006; 20-22: p<0.0001). For mice on the KEED (C+k and MP+k), neither effect (Genotype and Time) or their interaction reached significance, indicating C+k and MP+k mice did not differ after dietary intervention; however, the lack of a diet effect or diet-related interactions in the three-way ANOVA obfuscates the meaning of this lack of statistical significance.

#### Diet

For control mice (C-k and C+k), neither effect (Time and Diet) nor their interaction reached statistical significance. For MitoPark mice (MP-k and MP+k), there was a significant effect of time (F_(1,10)_=14.68, p=0.0033). Posthoc tests indicated that only MP-k mice deteriorated over time (MP-k: p=0.0180), but MitoPark groups did not differ from each other at either time point. Overall, these data suggest a potentially subtle effect of late diet intervention on delaying spontaneous movement deficits.

On the rotarod, before KEED intervention, there were no significant effects or interactions, although the effect of epoch was near significance (Two-way RM ANOVA, epoch: F_(1,23)_=3.939, p=0.0592). Sidak’s multiple comparisons tests revealed that although there were no differences between genotypes at either time point (p>0.05), only MitoPark mice differed in time spent on rotarod between early and mid-time points (Control: p=0.9600; MP: p=0.0252). After intervention, a three-way RM ANOVA indicated significant effects of time (F_(1,19)_=15.70, p=0.0008) and genotype (F_(1,19)_=33.84, p<0.0001), along with significant time x genotype (F_(1,19)_=5.121, p=0.0356) and genotype x diet (F_(1,19)_=6.916, p=0.0165) interactions. As before, we analyzed the data in separated two-way ANOVAs to characterize the significant effects and interactions.

#### Genotype

For SD mice (C-k and MP-k), there were main effects of time (F_(1,10)_ =8.994, p=0.0134) and genotype (F_(1,10)_=43.63, p<0.0001), but no interactions. Sidak’s post hoc tests suggest that only MP-k mice deteriorate in performance over time (C-k: p=0.4593; MP-k: p=0.0174), but MP-k mice perform worse than C-k mice at both time points (16-18: p=0.0005; 20-22: p<0.0001). For KEED mice (C+k and MP+k), there was an effect of time (F_(1,9)_=6.840, p=0.0280), but no genotype effect or time x genotype interaction. Like standard diet mice, Sidak’s tests indicated that C+k mice did not deteriorate over time (p=0.8887), but MP+k mice did (p=0.0159). Unlike standard diet mice, C+k and MP+k did not perform differently at the first time point post-intervention, though MP+k mice eventually developed deficits relative to C+k mice (16-18: p=0.8584; 20-22: p=0.0231). These results suggest that late intervention may delay development of rotarod deficits.

#### Diet

Control mice (C-k and C+k) did not differ across time or diet (p>0.05 for all). However, MitoPark mice (MP-k and MP+k) displayed significant main effects of time (F_(1,11)_=19.22, p=0.0011) and diet (F_(1,11)_=7.810, p=0.0174), but no significant interaction (p>0.05). Sidak’s tests indicate these effects were driven by deterioration across time in both groups (MP-k: p=0.0285; MP+k: p=0.0145), and MP+k mice performing better than MP-k mice immediately after KEED onset (16-18: p=0.0440) but equaled MP-k performance later (20-22: p=0.1326). These analyses provide further evidence that the KEED may delay rotarod motor deficits.

## Discussion

Ketone-ester enriched diets have been found to have beneficial effects for several conditions including diabetes^29, 30^, epilepsy,^18, 19^ Alzheimer’s Disease,^15^ heart disease^31^, and radiation damage^32^. Here, we demonstrate that a KEED could also preserve motor function in a PD model. The MitoPark model is notable for its progressive onset of motor deficits and dynamic response to L-DOPA treatment,^8, 9^ mirroring the progression of motor symptoms in human PD patients.^8, 33^ Consistent with previous studies, we found deficits in spontaneous (open field) and forced locomotion (rotarod)^34^ as well as dopamine neuron and striatal axon degeneration in MitoPark mice. However, MitoPark mice fed the KEED prior to motor deficit onset did not develop motor deficits through 20 weeks and had significantly increased dopamine release relative to MP-k controls. Notably, the preservation of dopamine was limited to evoked dopamine release as MP+k mice did not differ from MP-k mice in tonic tissue levels of striatal dopamine. In MP mice fed the KEED after the onset of motor deficits, there was no rescue of motor function, however, subsequent motor function degradation was delayed relative to MP-k mice.

Under normal conditions, the brain relies primarily on glucose metabolism as its primary energy source. However, temporal variations in the brain’s energy demand and supply require alternative fuel sources. For example, during periods of low glucose utilization or prolonged fasting, metabolism shifts to hepatic catabolism of triglycerides into fatty acids.^35^ However, since albumin-bound fatty acids are not permeable to the blood brain barrier, excess acetyl-CoA from β-oxidation is converted into ketone bodies: acetoacetate, acetone, and D-β-hydroxybutyrate (D-BHB). Recently, administration of exogenous D-BHB was shown to increase circulating ketone levels to ∼3.2 mM.^36-38^ D-BHB crosses the blood-brain barrier, cell membrane, and mitochondrial membrane using monocarboxylate transporters^39^, accumulating in the mitochondria, where it can be metabolized into acetyl-CoA through 3-hydroxybutyrate dehydrogenase 1.^40^ In this study, we utilized a D-BHB ketone ester enriched diet^22^ to increase blood ketone levels (Fig 1A).

We hypothesized that the KEED-mediated rescue of PD-like motor deficits in this study occur through two complementary biochemical mechanisms. Firstly, administration of D-BHB increases the availability of NADPH.^41^ Mechanistically, D-BHB has been shown to increase mitochondrial acetyl-CoA levels, which increases flux through the TCA cycle and increases citrate levels.^17, 42^ The increased citrate is oxidized to α-ketoglutarate in the mitochondria while NADP+ is reduced to NADPH.^43^ NADPH has many intracellular functions, one of which is the redox control of the dihydrobiopterin-tetrahydrobiopterin (BH2/BH4) couple. BH4 is a required cofactor for tyrosine hydroxylase^44^, and increasing its bioavailability would enhance catecholamine synthesis. Notably, PD patients have been shown to have reduced BH4 availability.^45^ Thus, increased BH4 levels may contribute to the ketone-mediated rescue of Parkinsonian symptoms by increasing dopamine synthesis. However, although evoked levels of dopamine were elevated in MP+k compared to MP-k, tonic striatal dopamine tissue levels did not differ significantly between groups, raising the possibility that the beneficial effect of the KEED is via enhanced vesicular packaging of dopamine. One possible mechanism supporting this would be through facilitation of VMAT2 activity, which is an active transporter, and D-BHB maintenance of ATP levels under hypoglycemic and increased mitochondrial dysfunction conditions^46^. Further work is required to test this hypothesis.

Dopaminergic neuron degeneration is a hallmark of PD. Contributing to this degeneration is oxidative stress and the decreased neuronal capacity to deal with reactive oxygen species (ROS).^47^ Reactive oxygen species were considered to be solely deleterious byproducts of metabolic reactions, but now are being increasingly appreciated for their role in intracellular signaling. For example, high levels of mitochondrial ROS can activate apoptosis/autophagy pathways.^48^ Several studies have demonstrated a D-BHB-mediated neuroprotective effect against oxidative stress.^46, 49-51^ Mechanistically, D-BHB functions as a direct antioxidant for hydroxyl radicals (^*^OH)^46^ and prevents mitochondrial production of ROS through NADH oxidation.^52^ As a histone deacetylase (HDAC) inhibitor, D-BHB may also transcriptionally activate oxidative stress resistance genes *FOXO3, MnSOD, CAT* and *MT2*.^51, 53^ However, we found that neither striatal terminals nor midbrain cell bodies were preserved in MP+k and MP-k groups. Thus, even though D-BHB may play a role in quenching ROS, this effect did not translate into whole cell neuroprotective effects. This may be in part due to conflicting evidence of D-BHB’s ability to be a HDAC inhibitor.^54^

An additional mechanism through which the KEED may benefit PD symptoms is by alleviating cellular respiration deficits that are prevalent in PD^7^ and a hallmark of the MitoPark model.^8, 11^ In vitro, D-BHB promotes increased ATP production.^17, 40, 46^ Furthermore, electron transport chain complex I dysfunction has been found to be a key driver of Parkinson’s pathogenesis.^55-58^ Many complex I inhibitors, such as 6-OHDA, MPTP, paraquat, and rotenone, cause dopaminergic degeneration and Parkinsonian symptoms.^59^ The catabolism of D-BHB produces a 1.5 fold increase in succinate^42^, a substrate for complex II, and is thus able to circumvent complex I dysfunction.

In addition to this study, several other studies have investigated the effects of other treatments in the MitoPark model. Langley, Ghosh ^60^ demonstrated the ability of a novel mitochondria-targeted antioxidant, Mito-Apocynin, to alleviate mitochondrial dysfunction, neurodegeneration, and behavioral deficits in MitoPark mice.^60^ Similarly, a recent study demonstrated the ability of quercetin, a plant flavonoid, to alleviate these deficits by improving mitochondrial bioenergetics.^61^ Furthermore, exercise has been shown to improve motor function in MitoPark mice due to increased aerobic respiration.^62^ These results, combined with those of the present study, indicate that restoring mitochondrial function may be a promising therapeutic intervention for PD patients. However, it is worth noting that the MitoPark model is specifically engineered to induce mitochondrial dysfunction,^8, 11^ and future studies should confirm the efficacy of mitochondrial-based interventions in other progressive models of PD, such as the α-synuclein overexpression model.^63^

In summary, the present study establishes the potential for the use of ketone esters in the treatment of Parkinson’s disease. MitoPark mice fed a KEED had preserved motor function and striatal dopamine release relative to MitoPark mice fed the standard diet. Although we were unable to determine a specific mechanism for this rescue, it is possible that by enhancing/preserving mitochondrial bioenergetics in MitoPark mice fed the KEED, dopamine synthesis and vesicular packaging were preserved and accounts for the amelioration of motor deficits and dopamine release deficits. Future studies will assess this possibility in MitoPark and other PD mouse models Overall, our study demonstrates a potential adjunct therapy for treating PD-related symptomology.

## Supporting information

Supplemental Methods & Results

## Acknowledgements

We are grateful to Gabriel Loewinger, Dr. Huaibin Cai, and William Curtis for helpful discussions about the data and to Ethan Chau for expert technical assistance.

## Dedications

This paper is dedicated to the memory of Dr. Richard “Bud” Veech whose belief in the broad therapeutic benefits of a ketone ester enriched diet inspired this study.

## Author Contributions

Designed Research: MID, RLV, AGS, DML

Contributed Essential Reagents: MTK

Performed Research: VRM, JAN, AGS, RJP

Analyzed Data: JAN, AGS, RJP

Prepared Manuscript: VRM, JAN, AGS, DML

## Data Availability Statement

The data that support the findings of this study are available from the corresponding authors upon reasonable request.

