## Supplemental Methods & Results for "Ketone ester-enriched diet ameliorates motor and dopamine release deficits in MitoPark mice"

Animals

All experiments and animal care were performed in compliance with the National Institutes of Health Care and Use of Animals guidelines and approved by the Institutional Animal Care and Use Committee of the National Institute on Alcohol Abuse and Alcoholism. MitoPark mice were generated by crossing mice with a loxP-flanked Tfam gene (Tfam^fl/fl^, Jackson Laboratory, strain #026123) with mice expressing Cre under the control of the DAT promoter (DAT-IRES-Cre, Jackson Laboratory, strain #006660) bred in-house on a C57BL6 background to selectively knockout Tfam in dopamine neurons.^1-3^ The mice used in this study were all homozygous for Tfam^LoxP^ and either heterozygous (MitoPark) or wild type (Control) expression of Cre recombinase. Mice were housed on a 12 h light/dark cycle on ventilated racks in a temperature- and humidity-controlled room with ad libitum access to food and water and no more than four mice per cage.

Fast-scan cyclic voltammetry

At 5, 10, or 20 weeks of age, mice were anesthetized with isoflurane and rapidly decapitated. Brains were extracted and immersed in ice-cold, carbogen-saturated (95% O_2_/5% CO_2_), cutting ACSF containing the following (in mM): Sucrose (194), NaCl (30), KCl (4.5), NaHCO_3_ (26), NaH_2_PO_4_ (1.2), dextrose (10), and MgCl_2_ (1). Coronal sections (300μm) spanning the striatum were prepared as previously described^4^ and incubated for 1 h before experiments in carbogen-saturated voltammetry recording ACSF (pH 7.4) containing (in mM): NaCl (126), KCl (2.5), NaHCO_3_ (25), NaH_2_PO_4_ (1.2), dextrose (10), HEPES (20), CaCl_2_ (2.4), MgCl_2_ (1.2), and L-ascorbic acid (0.4). Monophasic electrical pulses were generated with a DS3 Constant Current Stimulator (Digitimer, Ft. Lauderdale, FL) and delivered through a twisted, bipolar, stainless steel stimulating electrode (Plastics One, Roanoke, VA). The distance between the two poles of the stimulating electrode was adjusted to ~500μm. Carbon fiber electrodes were made as previously described^5, 6^ and cut to 150-300μm. The carbon fiber electrode potential was linearly scanned as a triangle waveform from −0.4 to 1.2 and back to −0.4V at 400V/s. Cyclic voltammograms were collected at 10 Hz using a Chem-Clamp (Dagan Corporation, Minneapolis, MN) and DEMON Voltammetry and Analysis software.^7^ After achieving stable baseline measurements, slices were stimulated every three minutes at various intensities (50, 100, 200, 400, 600, and 800µA, two stimulations per intensity) for 1 millisecond to evoke dopamine release and obtain an input-output curve. Extracellular dopamine concentrations were determined by post hoc calibrations against a 1 μM dopamine solution (Sigma-Aldrich, St. Louis, MO). All KEED mice used for these experiments began the diet at 4 weeks.

Immunofluorescence

Mice were anesthetized with pentobarbital and transcardially perfused with PBS followed by a 4% formaldehyde/PBS solution. The brains were then extracted and post-ﬁxed overnight before being stored in PBS for later sectioning. Fifty-micron thick sections were washed in PBS-T (0.2% Triton X-100), washed for 2 x 10 minutes in 0.5% NaBH4 in PBS to decrease autofluorescence, re-washed in PBS-T, then blocked for two hours with 5% BSA in PBS-T. Sections were then incubated overnight at 4°C in a primary antibody/0.5% BSA/PBS-T solution of rabbit anti-TH (1:1000, Invitrogen, Carlsbad, CA, 701949). Sections were then washed six times in PBS-T before incubating in secondary antibody/0.5% BSA/PBS-T solution of Alexa568 donkey anti-rabbit (1:2000) for two hours. Following six washes in PBS, sections were mounted onto subbed slides, cover-slipped with Fluoromount G containing DAPI (Electron Microscopy Sciences, Hatfield, PA), and imaged with a Zeiss AxioZoom microscope equipped with an Axiocam MR camera with ZEN software suite (Zeiss, White Plains, NY). Epifluorescence filter sets were standard for DAPI (350 excitation, 400 long pass beam splitter, 460/50 emission), and AF568/Texas red (560/40 ex, 585lp BS, 630/75 em). For comparative terminal field intensity measurements in the striatum, images were captured at 32x magnification, and exposure time was set to 50ms for the DAPI channel and 450ms for the TxRed channel. In the midbrain, images were captured at 125x magnification and the exposure times were 10ms for the DAPI channel and 35ms for the TxRed channel. Images were quantified using Fiji/Image J. For striatal terminal intensity measurements, images were imported and calibrated, then background fluorescence was subtracted by drawing a ROI in the lateral ventricle (or outside the slice, if the section was too anterior) and using ImageJ’s Process->Subtract Background function. An ROI was then drawn around the dorsal striatum, and average intensity across the dorsal striatum was measured. To count cells, each midbrain image was divided into SNc and VTA sections, and cells were manually counted in FIJI. Individual cells were selected as distinct, nucleated, circular/ovular structures. Quantification and image processing was performed blinded to the treatment and genotype of the mice. All KEED mice used for these experiments began the diet at 4 weeks.

Biochemistry

Procedures for the determination of brain tissue dopamine and DOPAC by gas chromatography-mass spectrometry (GC-MS) are described. Analytes were analyzed as the tertiary butyl dimethylsilyl ether-ester derivatives (TBDMS) using GC‐MS in the electron impact mode and quantified using the ^2^H_3_‐acetate.

Approximately 10mg of frozen brain tissue extracted from the dorsal striatum was added to 80μl of a 3.6% perchloric acid solution and in 2 ml polypropylene screw capped tubes with 10-12 small glass beads. Samples were shaken on a Mini Bead Beater (Biopsec Products, Bartlesville, OK) for 30 seconds (2x) until samples were homogeneous. Samples were placed on ice until centrifugation at 4°C in a Sorvall benchtop centrifuge at a speed of 10.7 xg for 2 min. Sixty microliters of the upper layer was pipetted into clean tubes and neutralized with 9.5μl 3M KHCO_3_.

Ten microliters of the sample extracts were evaporated under a stream of nitrogen to dryness in 1.5mL sylinized screw capped glass vials. Additionally, 6μl of a dilution of the labeled internal standards (^2^H_4_-dopamine, ^2^H_5_- DOPAC, ^2^H_3_-L-DOPA obtained from Toronto Research Corp., Toronto Ontario) (2, 10, 2 respectively mg in 400ul water). Samples were immediately reacted with 6μl of the sylilating reagent, N-methyl-N-(tri-methylsilyl) trifluoroacetamide (MTBSTFA) with 1% tert-butyldimethylchlorosilane (TBDMCS) reagent and the bis-trimethylfluoro methyl silyl (BSTFA) (Pierce Chemical Co., Rockford, IL) in 12μl of acetonitrile and heated to 60°C for 5 min.

Samples were analyzed on an Agilent 5973 quadrupole GC‐MS (Agilent, Wilmington, DE). One μl of the sample solutions was injected onto a 250μm × 30m capillary DB‐1 (Agilent) column in the splitless injection mode using helium as the carrier gas with a flow rate of 1.5 mL/min. The injector temperature was set at 250°C and the transfer line at 280°C. The GC oven temperature was programmed from 40 to 325°C at 15°C/min. The mass spectrometer was operated in the electron impact mode (70 eV) and the quadrupole mass analyzer scanned for ions which corresponded to a loss of 15 mass units (‐CH_3_) from the molecular ion of the derivative and its corresponding ^2^H‐labeled internal standard using selected ion monitoring mode. The ratios of the ions m/z 354/358 (dopamine retention time (RT), 13.1 min) (384/389 DOPAC, RT 12.85 min) and 470/473 (L-DOPA RT 15.2 min) were used to quantify the quantity of each analyte.

**Detailed Results**

*Chronic KEED elevates blood ketone ester concentration in a dose dependent manner*

We tested KEEDs with 16%, 32%, and 50% of carbohydrate calories replaced by BHB, alongside standard rodent chow and a control diet (Fig 1A). To verify that the KEED elevated blood ketone levels and determine the most appropriate diet concentration for further experiments, we compared blood concentration of ketones at 0, 2, 9, 16, 23, and 30 days after beginning a KE or control diet in C57BL6J mice. Due to severe weight loss (~33% after 16 days), mice on the 50% KEED were taken off the diet and removed from subsequent analysis. Two-way repeated measures ANOVA revealed significant effects of diet (F_(4,15)_=3.446, p=0.0346) and day (F_(1.831,27.47)_=27.03, p<0.0001), along with a significant interaction (F_(8,30)_=16.08, p<0.0001). Post-hoc tests indicated that mice on control diets did not change weight across time. Furthermore, although mice on the 32% KEED lost a significant amount of weight (Tukey’s tests, day 0 vs. 16, p=0.0063; day 0 vs. day 30, p=0.0180), mice on the 16% KEED did not lose weight (Tukey’s tests, 0 vs. 16, p=0.0866; 0 vs. 30, p=0.1448, Fig 1B). Two-way repeated measures mixed-effects modeling between the remaining groups’ blood ketone levels indicated significant effects of day (F_(2.016,56.04)_=10.33, p=0.0001) and diet (F_(4,31)_=84.10, p<0.0001), along with a significant interaction (F_(20,139)_=6.342, p<0.0001). Thus, the 16% and 32% KEEDs successfully increase blood ketone levels, but only the 16% KEED allowed mice to maintain initial weights and was thus used for all subsequent experiments.

*KEED ameliorates spontaneous locomotion deficits in the open field*

From weeks 6-20, MitoPark or control mice fed a KEED or standard diet (C-k, C+k, MP-k, MP+k) were tested weekly in the open field (to assess spontaneous locomotion) and on the accelerating rotarod (to assess forced locomotion). Behavioral data were binned into three epochs: Early (weeks 6-10), Mid (11-15), and Late (16-20). In the open field, mixed-effects three-factor mixed modeling analyses indicated significant main effects of week (F_(2,123)_=8.097, p=0.0005) and genotype (F_(1,64)_=15.92, p=0.0002), but not diet (F_(1,64)_=0.0001005, p=0.9920). Furthermore, there were significant week x genotype (F_(2,123)_=20.76, p<0.0001) and genotype x diet (F_(1,64)_=10.61 p=0.0006) interactions, and a significant week x genotype x diet interaction (F_(2,123)_=4.291, p=0.0158). To further investigate simple main effects, we performed two-way mixed-effects modeling across genotype and diet factors.

**Diet:** In control mice, there were significant effects of week (F_(1.457,48.82)_=4.591, p=0.0242), and diet (F_(1,35)_=5.081, p=0.0306), but no significant interaction (F_(2,67)_=2.290, p=0.1092). The significant effect of week was driven solely by an increased average velocity between Early and Mid epochs (p=0.0162). The significant effect of diet was driven by the C-k mice having a significantly higher average velocity than C+k mice in the Late epoch (p=0.0370). In MitoPark mice, there were significant effects of week (F_(1.762,49.33)_=31.78, p<0.0001) and diet (F_(1,29)_=5.872, p=0.0219), and a trend towards a significant interaction (F_(2,56)_=3.134, p=0.0513). The effect of week was driven by a decrease in average velocity across epochs (p<0.002 for all). The effect of diet was driven by the MP+k mice moving significantly more than the MP-k diet in Late weeks (p=0.0012). Furthermore, although MP-k mice significantly decreased average velocity across epochs (Early vs. Mid, p=0.0129; Mid vs. Late, p=0.0002; Early vs. Late, p<0.0001), MP+k mice only showed significant deficits when comparing Early and Late epochs (Early vs. Mid, p=0.1322, Mid vs. Late, p=0.4888, Early vs. Late, p=0.0004). Combined, these data suggest that MitoPark mice fed a KEED develop delayed deficits in spontaneous locomotion relative to MitoPark mice fed a standard diet.

**Genotype:** In mice fed a standard diet, there was a significant effect of week (F_(1.435,48.80)_=4.074, p<0.0351) and genotype (F_(1,36)_=22.62, p<0.0001), and a significant interaction (F_(2,68)_=16.70, p<0.0001). C-k mice displayed slower average velocity only on Early vs. Mid (p=0.0475), but not Early vs. Late (p=0.3758) or Mid vs. Late (0.4864) epochs. In contrast, average velocity in MP-k mice progressively declined across time points (Early vs. Mid, p=0.0129; Mid vs. Late, p=0.0002, Early vs. Late, p<0.0001). Furthermore, MP-k mice had significant spontaneous movement deficits at both Mid (p<0.0001) and Late (p<0.0001), but not Early (p=0.8130) time points, compared to C-k. In mice fed the KEED, there was a significant effect of week (F_(1.605,44.14)_=7.074, p=0.0040) and a week x genotype interaction (F_(2,55)_=7.045, p=0.0019). However, there was no effect of genotype (F_(1,28)_=0.4066, p=0.5289). The effect of week was driven by the MP+k mice, whose average velocity significantly declined between Early and Late epochs (p=0.0004) but not Early vs. Mid (p=0.1322) or Mid vs. Late (0.4888). C+k mice did not change average velocity across time (Early vs. Mid, p=0.4182; Mid vs. Late, 0.2088; Early vs. Late, 0.8924). Overall, the presence of distinct effects across genotypes in standard diet but not KEED mice further supports a beneficial effect of the KEED.

*KEED ameliorates forced locomotion deficits on the accelerating rotarod*

To assess forced locomotion, motor coordination, and balance, MitoPark and Control mice fed either the KEED or standard home chow were run on an accelerating rotarod test every week from 6 to 20 weeks of age. Mixed-effects three-factor mixed modeling analyses revealed significant main effects of week (F_(2,113)_=59.53, p<0.0001) and genotype (F_(1,59)_=12.35, p=0.0009), but not diet (F_(1,59)_=1.424, p=0.2375)**.** Furthermore, there were significant week x genotype (F_(2,113)_=18.64, p<0.0001) and week x diet (F_(2,113)_=0.5810, p=0.0040) interactions**,** and a significant week x genotype x diet interaction (F_(2,113)_=4.956, p=0.0086). To further investigate simple main effects, we performed two-way mixed-effects modeling, with mice split by genotype or diet factors.

**Diet:** In control mice, there was a significant effect of week (F_(1.863,54.96)_=5.745, p=0.0064), but no effect of diet (F_(1,30)_=0.01956, p=0.8897) or interaction (F_(2,59)_=0.05925, p=0.9425). Tukey’s multiple comparisons tests indicated the significant effect of week was driven by a decreased latency to fall at late weeks points relative to early weeks (p=0.0098) across diets. In MitoPark mice, there was a significant effect of week (F_(1.895,51.16)_=75.88, p<0.0001) and a significant week x diet interaction (F_(2,54)_=11.22, p<0.0001), but no significant effect of diet alone (F_(1,29)_=3.3702, p=0.0642). Both MitoPark groups showed progressive decline across time (MP-k: all comparisons p<0.0001; MP+k: Early vs. Mid, p=0.0264, Mid vs. Late, p=0.0166, Early vs. Late, p=0.0007). However, the MP+k mice had significantly longer latency to fall in Late weeks compared to the MP-k mice (p=0.0014), indicating the KEED lessens the degree of forced locomotor deficits. Overall, these data indicate the KEED may improve forced locomotion in MitoPark mice.

**Genotype:** In mice fed a standard diet, there was a significant effect of week (F_(1.886, 55.65)_=50.10, p<0.0001) and genotype (F_(1,31)_=12.18, p=0.0015), and a significant interaction (F_(2,69)_=21.05, p<0.0001). Multiple comparisons tests indicated that C-k mice did not change performance across weeks (Early vs. Mid, p=0.6286; Mid vs. Late, p=0.2366, Early vs. Late, p=0.2366), but in MP-k mice, latency to fall decreased across weeks (all p<0.0001). Although latency to fall in MP-k mice declined across epochs, these mice only differed from the Control - ketone mice at Late weeks (p<0.0001). In mice fed the KEED, there was a significant effect of week (F_(1.889,51.01)_=14.79, p<0.0001). However, there was no effect of genotype (F_(1,28)_=2.238, p=0.1459) and no significant week x genotype interaction (F_(2,54)_=2.259, p=0.1143). Tukey’s multiple comparisons tests indicate that the effect of week is driven by significant, or near-significant, differences between all time points (Early vs. Mid, p=0.0081; Mid vs. Late, p=0.0580; Early vs. Late, p=0.0003). These analyses also support the ameliorative effect of the KEED on forced movement and motor coordination.

*KEED ameliorates evoked dopamine release deficits*

To determine the time course of dopamine release degradation and the effect of the KEED, we measured evoked dopamine release in striatal slices using FSCV. We compared control and MitoPark mice fed standard diets at 5, 10, and 20 weeks of age. At 5 weeks, control and MitoPark mice did not differ in evoked dopamine release (genotype: F_(1,16)_=0.1154, p=0.7385; genotype x intensity interaction: F_(5,80)_=0.9360, p=0.4624), along with a significant main effect of stimulation intensity. At 10 weeks, however, MitoPark mice had drastically lower dopamine release overall (significant main effect of genotype, F_(1,18)_=8.902, p=0.0080; significant main effect of intensity, F_(1.185,21.33)_=22.44, p<0.0001; significant interaction, F_(5,90)_=3.032, p=0.0142). This effect was driven by decreased dopamine release at 200, 400, 600, and 800 µA (Sidak’s multiple comparisons tests, p<0.05 for all).

At 20 weeks, we also examined the effect of the KEED on dopamine release. A three-way ANOVA revealed significant effects of intensity (F_(1.524,47.23)_=55.02, p<0.0001) and genotype (F_(1,31)_=57.27, p<0.0001), along with a significant genotype x intensity interaction (F_(5,155)_=37.97, p<0.0001). All other effects and interactions were not significant. Simple main effects analysis indicated that dopamine release in C-k and C+k mice did not differ (diet: F_(1,16)_=0.08613, p=0.7729; intensity x diet: F_(5,80)_=0.3030, p=0.9097). However, dopamine release significantly differed between MP-k and MP+k groups, indicated by a significant effect of diet (F_(1,15)_=18.74, p=0.0006) and significant intensity x diet interaction (F_(5,75)_=19.43, p<0.0001). Post-hoc tests indicated these effects were driven by significant or near-significant increases in dopamine release in the MP+k mice at 600 (p=0.0483) and 800 (p=0.0579) µA. These results suggest that the KEED boosts dopamine release at 20 weeks in MitoPark mice.

*KEED does not increase cell survival, striatal TH+ axons, or tissue dopamine content*

To determine if the KEED rescue of dopamine release was due to preservation of dopamine neurons or axons, we performed immunohistochemistry for tyrosine hydroxylase (TH) (Fig 4A). Across the anterior-posterior gradient of the dorsal striatum, MitoPark mice had significantly decreased TH+ immunoreactivity compared to controls (three-way repeated measures mixed-effects analysis, significant main effect of genotype, F_(1,12)_=435.0, p<0.0001), indicating dopamine axon degeneration regardless of diet (no significant main effect of diet, F_(1,12)_=0.5335, p=0.4791; no significant effect of AP coordinate, F(_1.595,18.61)_=3.068, p=0.0799)) (Fig 4B). Similarly, counting of TH+ cells in the SNc and VTA indicated a genotype- but not diet-dependent decrease in cell count (three-way repeated measures ANOVA, significant main effect of genotype, F_(1,12)_=19.60, p=0.0008; no significant main effect of diet, F_(1,12)_=0.02963, p=0.8662) (Fig 4C). GC-MS analyses of flash-frozen striatal tissue indicated that, although dopamine and DOPAC content was significantly diminished in MitoPark mice (two-way ANOVAs, dopamine: significant main effect of genotype, F_(1,31)_=245.1, p<0.0001; DOPAC: significant main effect of genotype, F_(1,31)_=81.38, p<0.0001), but the KEED did not increase content of either compound (two-way ANOVAs, dopamine: no significant main effect of diet, F_(1,31)_=0.1231, p=0.7280; DOPAC: no significant main effect of diet, F_(1,31)_=1.420, p=0.2425) (Fig 4D-E). Sidak’s multiple comparisons tests confirmed that MitoPark mice, regardless of diet, had similar dopamine (p=0.7575) and DOPAC (p=0.9998). Across groups, mice did not differ in DOPAC/DA ratio (two-way ANOVA, diet: F_(1,31)_=1.688, p=0.2034; genotype: F_(1,31)_=2.198, p=0.1483; interaction: F_(1,31)_=1.326, p=0.2583), thought to be a correlate of MAO activity and levels of oxidative stress.^29^ Thus, although the KEED is able to rescue or ameliorate motor deficits and evoked dopamine release, it does not seem to slow degeneration or boost dopamine levels in the striatum.

*Late intervention delays but does not rescue motor deficits*

Although our KEED intervention successfully prevented the development of motor deficits at our endpoint of 20 weeks, a caveat of these experiments is that we began KEED administration at 4 weeks—well before any deficits emerged. This treatment schedule, however, is not comparable to treatment of PD in humans, as intervention typically begins after symptoms emerge. To this end, we tested whether a delayed start on the KEED, after the emergence of deficits, would delay or even rescue symptom progression. Mice began rotarod and open field testing at 8 weeks and KEED administration began at 14 weeks (after the emergence of both open field and rotarod deficits in MP-k mice, Fig 2), the endpoint was extended to 22 weeks to better capture the potential effects of KEED intervention (Fig 5A). As with the behavioral data in the original experiments, we binned data into equal epochs (8-10, 12-14, 16-18, and 20-22 weeks).

Before dietary intervention (first two epochs) in the open field, we found a significant effect of time (two-way RM ANOVA, F_(1,20)_=47.95, p<0.0001) and a significant time x genotype interaction (F_(1,20)_=18.12, p=0.0004), but no effect of genotype alone (F_(1,20)_=2.886, p=0.1049). Sidak’s multiple comparisons tests indicated that MitoPark, but not control mice decreased movement from early to mid time points (MitoPark: p<0.0001; Control: p=0.1645), and that at the mid but not early time point, MitoPark mice moved significantly less than controls (early: p=0.9260; mid: p=0.0031). After dietary intervention (final two epochs), we observed a significant effect of genotype (Three-way RM ANOVA, F_(1,18)_=20.24, p=0.0003), and significant time x genotype interaction (F_(1,18)_=14.03, p=0.0015). All other effects and interactions were not significant (p>0.05). To further investigate the effects that reached significance, we analyzed the data across genotype and diet factors with two-way RM ANOVAs.

**Genotype:** For mice on the standard diet (C-k and MP-k), there was a significant effect of genotype (F_(1,10)_=38.68, p<0.0001) and time x genotype interaction (F_(1,10)_=12.94, p=0.0049). Sidak’s multiple comparisons tests revel that this effect is due to MP-k (but not C-k) mice moving less over time (MP-k: p=0.0031; C-k: p=0.5220), and MitoPark mice moving less at both post-diet time points (16-18: p=0.0006; 20-22: p<0.0001). For mice on the ketone diet (C+k and MP+k), neither effect (Genotype x Time) or their interaction reached significance, indicating C+k and MP+k mice did not differ after dietary intervention; however, the lack of a diet effect or diet-related interactions in the three-way ANOVA obfuscates the meaning of this lack of significance.

**Diet:** For control mice (C-k and C+k), neither effect (Time and Diet) nor their interaction reached statistical significance (p>0.05). For MitoPark mice (MP-k and MP+k), there was a significant effect of time (F_(1,10)_=14.68, p=0.0033), but diet and interaction were not significant. Posthoc tests indicated that only MP-k mice deteriorated over time (MP-k: p=0.0180; MP+k: p=0.0885), but MitoPark mice did not differ from each other at either time point. Overall, these data suggest a potentially subtle, but inconclusive effect of late diet intervention on delaying spontaneous movement deficits.

Before intervention on the rotarod, there were no significant main effects or interactions, although the effect of epoch was near significance (Two-way RM ANOVA, epoch: F_(1,23)_=3.939, p=0.0592; genotype: F_(1,23)_=1.155, p=0.2936; interaction: F_(1,23)_=2.564, p=0.1230). Sidak’s multiple comparisons tests revealed that although there were no differences between genotypes at either time point (p>0.05), only MitoPark mice differed in time spent on rotarod between early and mid time points (Control: p=0.9600; MP: p=0.0252). After intervention, a three-way RM ANOVA indicated significant effects of time (F_(1,19)_=15.70, p=0.0008) and genotype (F_(1,19)_=33.84, p<0.0001), along with significant time x genotype (F_(1,19)_=5.121, p=0.0356) and genotype x diet (F_(1,19)_=6.916, p=0.0165) interactions. As with open field, we then analyzed the data in separate two-way ANOVAs to further characterize the significant differences driving the significant effects and interactions.

**Genotype:** For mice fed a standard diet (C-k and MP-k), there were significant main effects of time (F_(1,10)_ =8.994, p=0.0134) and genotype (F_(1,10)_=43.63, p<0.0001), but no interaction (p>0.05). Sidak’s post hoc tests suggest that, as in open field, only MP-k mice deteriorate in performance over time (C-k: p=0.4593; MP-k: p=0.0174), but MP-k mice perform worse than C-k mice at both time points (16-18: p=0.0005; 20-22: p<0.0001). For mice fed the KEED (C+k and MP+k), there was a significant main effect of time (F_(1,9)_=6.840, p=0.0280), but genotype and time x genotype interaction were not significant. As with the standard diet mice, Sidak’s tests indicated that C+k mice did not deteriorate over time (p=0.8887), but MP+k mice did (p=0.0159). Unlike standard diet mice, C+k and MP+k did not perform differently at the first time point post-intervention, though MP+k mice eventually developed deficits relative to C+k mice (16-18: p=0.8584; 20-22: p=0.0231). These results suggest that late intervention may delay rotarod deficits.

**Diet:** Control mice (C-k and C+k) did not differ across time or diet (p>0.05 for all). However, MitoPark mice (MP-k and MP+k) displayed significant main effects of time (F_(1,11)_=19.22, p=0.0011) and diet (F_(1,11)_=7.810, p=0.0174), but no significant interaction. Sidak’s tests indicate these effects were driven by deterioration across time in both groups (MP-k: p=0.0285; MP+k: p=0.0145), and initial improved performance in MP+k after dietary intervention, but equaled MP-k performance after time (16-18: p=0.0440; 20-22: p=0.1326). These analyses provide further evidence that KEED delays development of deficits on the rotarod, and overall suggest that later intervention with the KEED may delay (but not rescue) motor deficits.

Supplemental Materials and Methods References
